# On use of tertiary structure characters in hidden Markov models for protein fold prediction

**DOI:** 10.1101/2024.04.08.588419

**Authors:** Ashar J. Malik, Caroline Puente-Lelievre, Nicholas Matzke, David B. Ascher

**Author notes:** Correspondence to Ashar J. Malik, David B. Ascher.

## Abstract

While advances in protein structure prediction have opened up insights into arcane proteins, weak sequence homology makes functional characterisation challenging. To overcome this challenge, we use structure-based hidden Markov models of groupings in SCOP, CATH and ECOD to predict folds in proteins and thereby infer function. Conservation of structure and ability of hidden Markov models to detect remote signals make this a powerful resource for complete characterisation of arcane proteins.

## Main text

Metagenomics has powered the search for enzymes that underpin industrial biotechnological applications, however their identification has primarily been limited to sequence similarity. The protein structure prediction revolution has opened up opportunities to explore understudied proteins that have limited sequence homology or functional characterisation, but untapped potential for biotechnological applications. Despite this, functional annotation of arcane proteins remains a challenge.

In the current era, marked by highly accurate protein structure prediction techniques [1, 2], a significant proportion of proteins in repositories like MGnify90 [3], Swiss-Prot [4], and Uniprot [5] have now been predicted [1, 2]. This advancement is shifting the paradigm from the “dark proteome” where proteins lack known structure and function [6], to the “gray proteome”, characterised by proteins with predicted structures but still undiscovered functions.

Conventionally the process of function determination involves comparing the sequence of the novel protein to those for which function is already known [7]. However, advancements in sequencing technologies [8] has discovered many “arcane” proteins, for which comparisons to characterised sequences frequently produces results in or beyond the “twilight-zone” [9], a theoretical limit in which a signal of evolutionary relatedness cannot be readily discerned from noise. In contrast to this, protein structures are known to be far better conserved over evolutionary time-scales, compared to the underlying sequence [10]. A structure-based comparison therefore permits deeper relationships to be uncovered. Three popular hierarchical databases, namely SCOP [11], CATH [12] and ECOD [13] leverage structural robustness and gather structures based on shared structural and functional similarity into hierarchical groups, where gathered proteins share similar folds. Within these groupings sequence similarity is often observed to be low. Several structure comparison tools can therefore detect if a novel protein for which a structure has just been experimentally resolved or predicted has sufficient structural overlap with proteins characterised within these databases. A high degree of structural overlap is clearly suggestive of grouping while a poor overlap results in failed characterisation.

In cases where structure-based comparison results in a low score, a machine learning predictor may prove useful. Such a predictor would be trained on characteristic features of each grouping and use the trained model to predict groupings of a query protein. These models have been employed in the past both relying on sequence-based [14] and structure-based features [15] with limited success. Classification of structures into groups as undertaken by SCOP, CATH and ECOD is a non-trivial task. While for some groups in these databases clear boundaries may be present, for others boundaries may be ambiguous. A simple way to illustrate this is to compare structures within groups and across groups. This exercise reveals that on a scale of zero to one, with zero representing no similarity and one indicating identity, proteins can have low comparison scores (TM-scores, using Foldseek [16]) of “0.3” with their group neighbours and proteins across groups can have high scores of “0.7” (see Figures S1-S6). While addition of other metrics such as alignment length may provide more resolution, these scores alone indicate that any structure-based features extracted for the purpose of training of machine learning-based predictors may also overlap and consequently lead to poor training and inaccurate classification.

This work uses Hidden Markov Models (HMMs) to circumvent this. HMMs have been shown to detect remote homology [17] and as a result considerable effort has been placed into developing resources like gene3D [18] and Superfamily [19] which can be used to obtain CATH and SCOP groupings for a query protein sequence. Although these efforts have increased sensitivity of sequence-based searches for group assignment in these respective databases, challenges remain in characterisation of deeply diverging proteins [20] as demonstrated later.

Traditionally HMM profiles of protein families are compared to a query amino acid sequence to determine if the HMM model can emit the query sequence. We employ HMMs in a similar capacity with the major difference of using protein structure data to generate HMMs instead of relying on amino acid sequences [21]. The models are then used to predict SCOP, CATH and ECOD groupings to generate functional insights. To generate structure-based HMMs, this work uses a relatively new protein structure comparison tool, Foldseek. Fast and accurate structural similarity determination, by Foldseek, is achieved by encoding the three dimensional (3D) structures into linear sequences of characters (3Di characters) prior to comparison. Each amino acid position is assigned one of 20 3Di characters which captures its immediate structural neighbourhood. We use this 3Di sequence of tertiary-interaction characters to build structure-based HMMs. This approach towards comparing protein structures has already demonstrated great promise [22]. By developing an HMM model which is based on 3Di characters to represent each of the proteins in a particular group, it is possible to check if a model can emit the query structure (also represented by its 3Di sequence) (see Figure 1 (b)).

**Figure 1:**
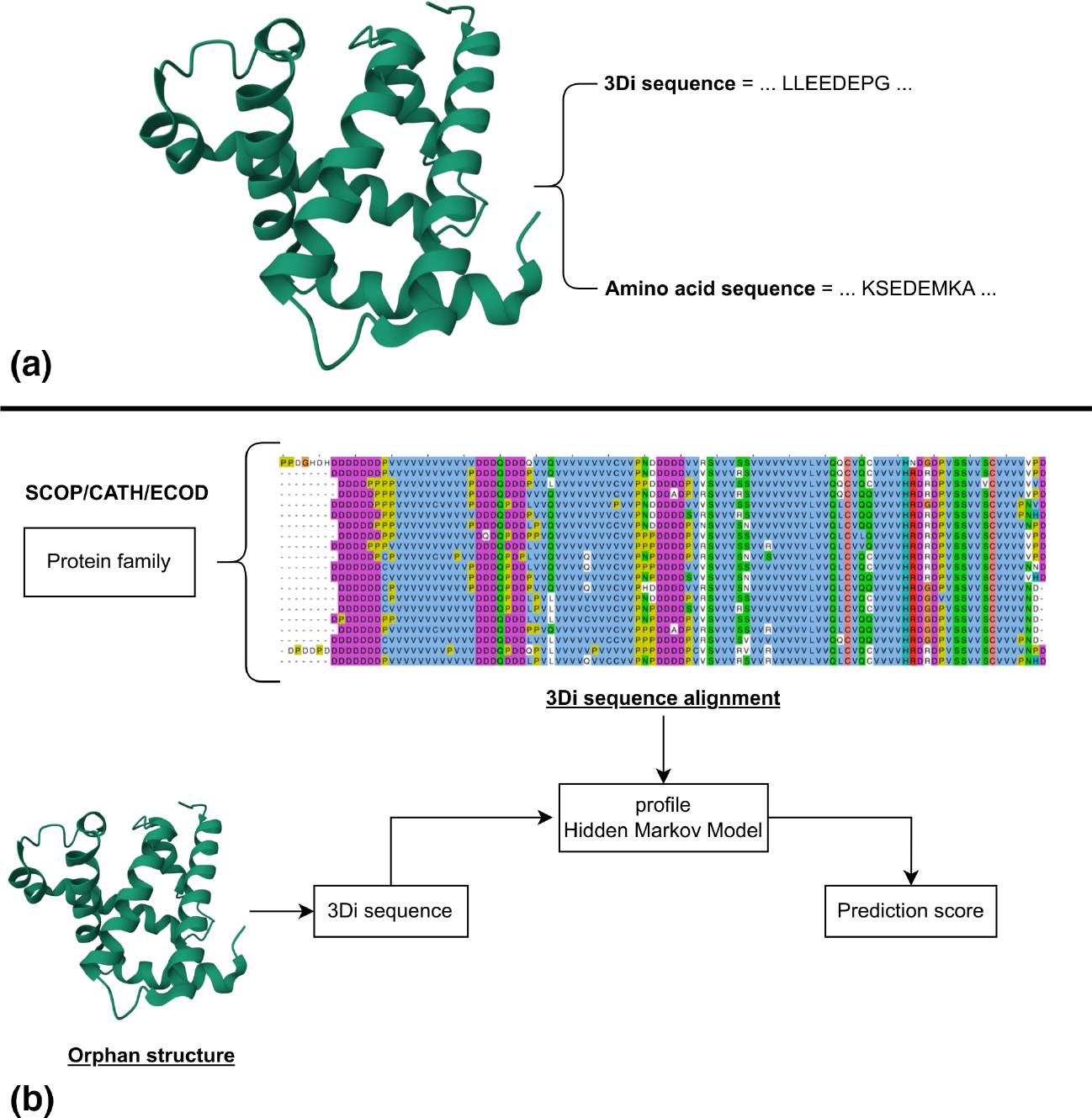
Utilisation of 3Di character-based hidden Markov models for fold prediction. (a) A cartoon representation of a protein structure is shown together with its representation in sequence-space, as a conventional amino acid sequence and the novel Foldseek generated 3Di sequence. (b) For any grouping in SCOP, CATH or ECOD, all member proteins are represented by their 3Di sequences, alignments are generated and converted to profile HMMs using HMMER. Upon querying, a protein structure is converted to its respective 3Di representation and scored against all SCOP, all CATH and all ECOD models and best scores are used to predict folds from each of the respective databases.

We generate over 5,600 HMMs for CATH, over 4,400 HMMs for SCOP and over 15,000 HMMs for ECOD, using HMMER [23], where each model represents its respective protein grouping in these databases. A web-based server (see Availability) is provided for users to input protein structures, which are then converted to their representative 3Di sequences and scored against all models for SCOP, CATH and ECOD respectively. The output provides a breakdown of the results per database along with a graphical output summarising the results.

Given all models can be tested to see emission probabilities against a query, we systematically investigated results to establish thresholds, which may assist with group assignment. While SCOP, CATH and ECOD each have four primary hierarchical levels, these are not equivalent which is why we use the generic term “groups” instead of terms like “class” and “family” to avoid confusion. For testing we compared protein pairs with increasing similarity (see Method), determined by their respective classifications. This analysis showed (see Figure 2 & Figures S7-S10; Supplementary sheet R1) that HMM emission scores indicating relatedness gradually increased as structural relatedness in proteins compared became higher. A leave-1-out strategy was developed where one protein was randomly removed from a group in the model building stage and then compared back to the built model in a validation step (see Figure S11; Supplementary sheet R2). This showed that on average the models generated high scores similar to scores obtained when proteins compared belonged to the same group. In another test we wanted to see the top-10 results of predicted groupings for proteins with known assignments. We compared a well distributed set comprising 1,000 structures to all HMMs and observed *>*98% classification accuracy across their respective databases, i.e., predicted groupings for a structure matched those assigned to it by SCOP, CATH and ECOD respectively (see Figure S12; Supplementary sheet R3).

**Figure 2:**
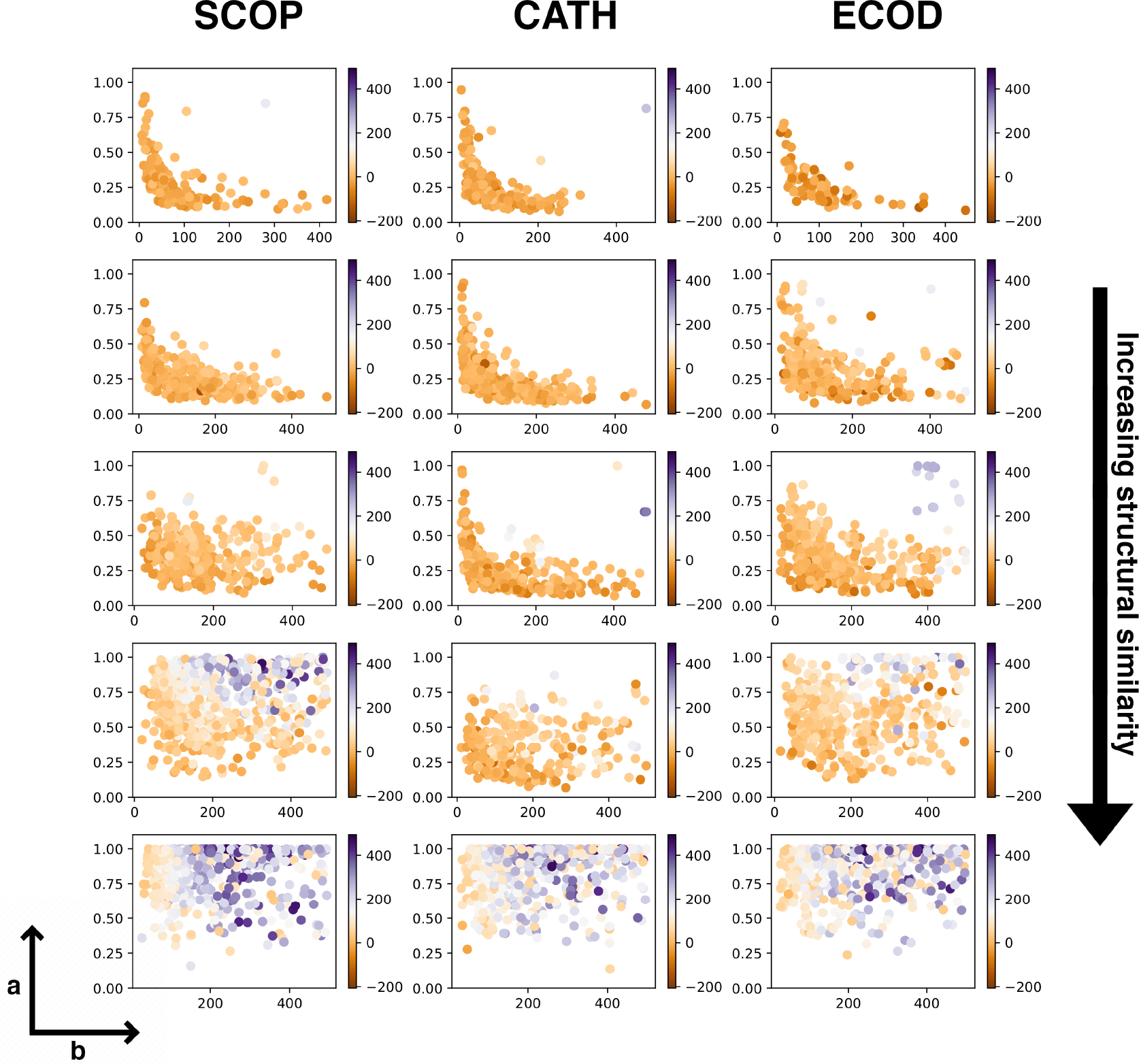
Trends in structure-based metrics with increasing structural similarity. The figure grid shows comparison of protein pairs for SCOP, CATH and ECOD respectively with structural similarity on (a) the vertical axis and (b) number of residues aligned in the structural alignment on the horizontal axis for all plots. For a given comparison colour represents the HMM scores. The data progressively shifts from the “twilight zone”-equivalent region (bottom-left region in the plot), characterised by a region below a decaying curve, to the safe region (top and top-right region in the plot). HMM scores obtained for these comparisons show a corresponding increase with increasing structural similarity. See Figures S7-S10 for more details.

For comparative statistics we included querying of amino acid sequences using HMMER against the Superfamily HMM database for SCOP assignments at the superfamily level (level 3 of SCOP hierarchy) and the gene3D database for CATH groupings at their respective homology level (level 4 of CATH hierarchy). For a randomly collected set of proteins (see Supplementary sheet R4), when true SCOP and CATH groupings were known (i.e., proteins had been classified by SCOP and CATH), we report an accuracy of 99% for SCOP and 94% for CATH when querying structure-based HMMs using 3Di sequences against 99% for SCOP and 98% for CATH using amino acid sequences against Superfamily and gene3D databases respectively. When true SCOP and CATH labels were unknown (i.e., proteins had not been classified by SCOP and CATH), we compared our results with those generated through sequence-based HMM querying and found that the two predictions were in agreement at 72% for SCOP and 82% for CATH.

In the cases where the predictions from structure-based and sequence-based HMMs were not in agreement, an explanation cannot be easily furnished. This work is based on the premise that structure is more conserved then sequence and therefore may lead to better grouping. An attempt to compare all amino acid sequences of proteins in a grouping back to the respective HMM of that group reveals that for some groups certain members score poorly. For the same members, structure-based comparison generated high scores. This is possible because while each protein in a group may have a distinct amino acid sequence which diverges on evolutionary time-scale, a 3Di sequence remains conserved as the structure remains similar and therefore is a stronger candidate for comparison purposes. So while the addition of amino acid sequence-based HMMs, to CATH and SCOP, through gene3D and Superfamily increase their respective abilities to find a grouping, these continue to struggle when diverged protein sequences are used. A case for this can be made using the example of Mitochondrial complex I, ND4L subunit from *Ovis aries*. A structure representing this protein has PDB ID 5lnk, chain K and carries the CATH assignment of 1.10.287.3510. The structure of this protein generates a high score (see Supplementary sheet R5) against the structure-based HMM for this group, however a sequence search against the same HMM in gene3D returns a low score with the web version of HMMER [24] providing no hits altogether. For SCOP, a case can be made using the example of Palmitoleoyl-protein carboxylesterase NOTUM from *Homo sapiens*. The structure PDB ID 4uz5, chain A, representing this protein carries the SCOP assignment of c.69.1.42. Like before this protein has a low score when querying using amino acid sequence but scores highly when compared to the 3Di sequence-based HMM for its respective group and as noted before, the web interface of HMMER does not return a result. While the above were simple cases which could be highlighted as the true SCOP and CATH assignments were known, the ESM Atlas alone holds more than 700 million predicted protein structures. A randomly selected example (ESM Magnify ID: MGYP003636274702) from that resource shows the same pattern, generating results only with this structure-based approach.

This resource presents an important advance as it combines the capacity of protein structures to hold evolutionary signals over longer time scales with the ability of HMMs to harvest deeper evolutionary relationships. Together, this approach can help move proteins, such as in the case of ESM Atlas, from the “gray proteome” to being fully characterised.

## Methods

### Protein structure and sequence data

All protein data (both amino acid sequence and structure) used in this work was obtained from RCSB PDB (rcsb.org), with the exception of the ESM structure carrying ID MGYP003636274702 which was obtained from the ESM Metagenomic Atlas.

VMD [25] was used to process structures which included extracting relevant chains from protein structures.

### SCOP, CATH and ECOD annotations

Complete SCOP (v2.08), CATH (v4.3.0) and ECOD (v288) assignments for downloaded from their respective online resources.

For SCOP, only the four primary classes characterised by “a”,”b”,”c” and “d” labels were retained for this work. For CATH, only three primary classes characterised by “1”,”2”,”3” labels were used. As ECOD follows a different hierarchical structure, all assignments were retained.

VMD was used to only extract portion of the structure from their respective PDB files which matched domain assignment in SCOP, CATH and ECOD. These ranges were obtained from the parseable database files obtained from their respective databases.

### What is a group?

The three popularly used structural characterisation databases, SCOP, CATH and ECOD each have primarily four levels of hierarchy each representative of similarity at a different scale. To avoid any confusion, this work uses the general term “group”.

We refer to the hierarchical levels as “H1”,”H2”,”H3”,”H4” so a typical assignment of a group would be denoted as “H1.H2.H3.H4”. This four-level assignment, for a query structure, is predicted in this work. An additional level of “H0” was created preceding “H1” for data analysis carried out in this work.

To generate statistics shown in figures (Figure 2 & Figures S7-S10; Supplementary sheet R1), both structures (using Foldseek) and sequences (using protein BLAST [26]) of protein pairs (P1, P2) were compared. The comparisons drew 1,000 pairs with increasing level of similarity. The first set drew 1,000 pairs at “H0”, meaning that both P1 and P2 had different “H1” classifications. This was followed by four additional runs at 1) “H1”, where P1 and P2 shared only the outer most assignments (e.g., P1: 1.30.70.100 and P2: 1.70.60.40), 2) “H1 & H2”, where P1 and P2 shared only the outer two assignments (e.g., P1: <u>2.30</u>.70.90 and P2: <u>2.30</u>.30.20), 3) “H1 & H2 & H3”, where P1 and P2 shared three assignments (e.g., P1: <u>3.30</u>.20.20 and P2: <u>3.30.20</u>.90) and finally 4) “H1 & H2 & H3 & H4”, where both P1 and P2 shared the same group (e.g., P1: <u>2.30.40.50</u> and P2: <u>2.30.40.50</u>).

For each pair six quantities were computed, namely structural similarity (TM-score), structural alignment length (Number of residues aligned in the structural alignment), sequence similarity (protein BLAST), sequence alignment length (Number of residues aligned in the sequence alignment) and two emission probabilities where P1 (3Di sequence) was compared to the HMM model to which P2 was assigned and where P2 (3Di sequence) was compared to the HMM model to which P1 was assigned. The two emission scores were averaged. Results are only shown where both sequence and structure alignment steps were successful (see Supplementary sheet R1)

### Superfamily and gene3D databases

To carry out sequence-based HMM scoring on a local high performance machine, these databases were downloaded from the target database page on EBI.

### Structure comparison and generation of 3Di characters

Protein structure comparison carried out in this work made use of the Foldseek structure comparison program. The 3Di sequences were also generated using Foldseek.

### HMM construction

All 3Di sequences were aligned using clustalw2 [27] using the scoring matrix provided in the original Foldseek publication to generate grouping specific multiple sequence alignments. The HMMER program was then used to construct HMM models for each group in SCOP, CATH and ECOD. In cases where only one member was present in the group, a duplicate entry of the member was created to take the member count of the group to two. This minimal data augmentation ensured HMM creation by HMMER.

### Web server

On querying, a structure is either provided by the user or obtained from RCSB PDB. Foldseek is used to obtain the respective 3Di sequence which is then compared to the HMMs developed in this work using HMMER in a database specific manner.

Foldseek is used to compare protein structures in a nearest neighbour (NN) search. The NN results show pre-assigned protein classification, allowing the users two modes from which to infer fold assignment. The Foldseek generated transformation matrix is applied to the structure using VMD and the resulting comparison is visualised in the web browser using the Mol^*\**^ [28] library.

The HMM search lists HMMER results along with all GO terms captured for all proteins in a particular group. RCSB saguaro protein feature viewer is used to show an interactive panel to visualise domains in the structure. The panel only shows the best scoring groups with the condition that the length of the predicted domain must comprise 40 residues; an arbitrary cutoff to avoid low scoring hits.

The server makes use of Flask, nginx, jQuery, html, and javascript.

## Supporting information

Supplemental Sheet 1

Supplemental Sheet 2

Supplemental Sheet 3

Supplemental Sheet 4

Supplemental Sheet 5

## Availability

A webserver is available here: https://biosig.lab.uq.edu.au/hmmpredict/

To allow for batch predictions, the HMM files can be requested from the authors.

## Notes

### Competing Interest Statement

The authors have declared no competing interest.

