## Supplemental PDF for "On use of tertiary structure characters in hidden Markov models for protein fold prediction"

### Supplementary material

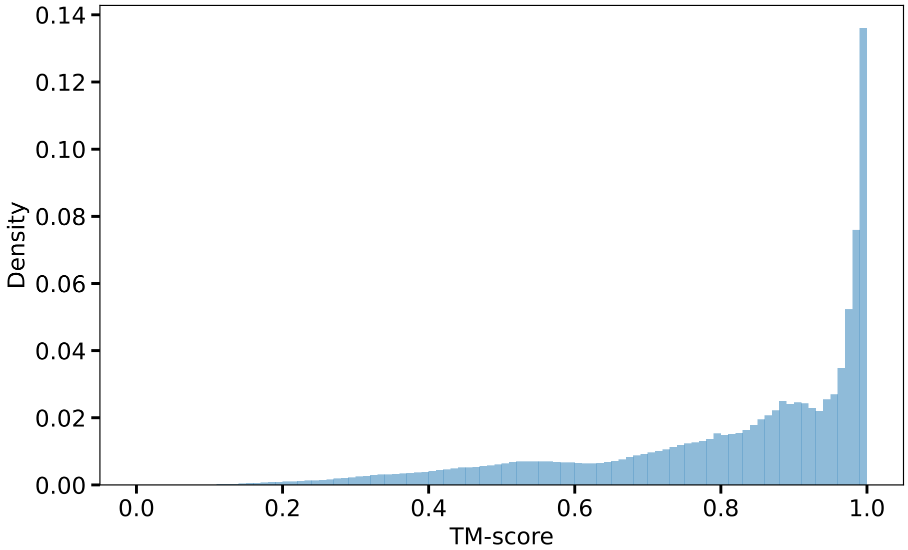


Figure S1: Distribution of pairwise structure comparison scores of proteins inside SCOP groups. Given a SCOP group with $N$ members, $N^2$ comparisons were attempted. This was repeated for all groups in SCOP. The figure shows distribution of structural similarity scores where comparisons were successful. The tail of the distribution extending towards 0 indicates low pairwise structural similarity between proteins in the same group.

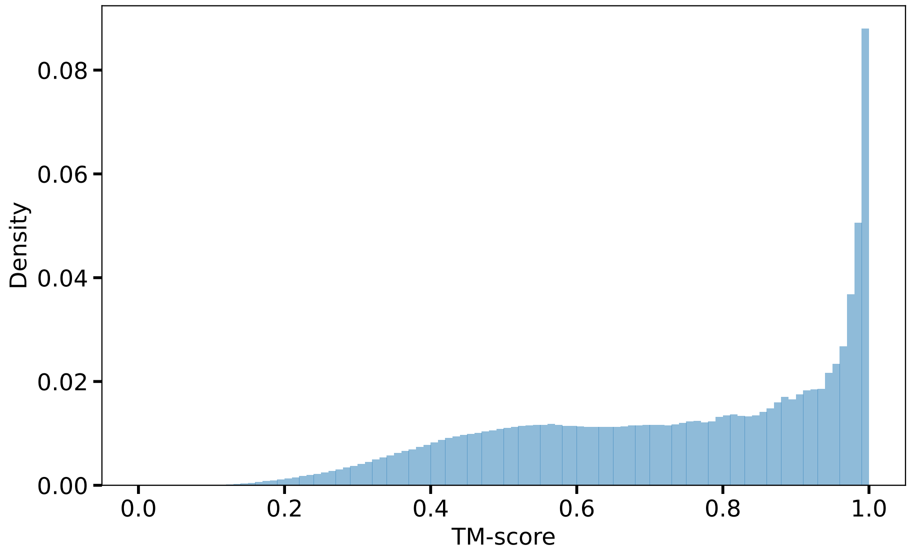


Figure S2: Distribution of pairwise structure comparison scores of proteins inside CATH groups. Given a CATH group with $N$ members, $N^2$ comparisons were attempted. This was repeated for all groups in CATH. The figure shows distribution of structural similarity scores where comparisons were successful. The tail of the distribution extending towards 0 indicates low pairwise structural similarity between proteins in the same group.

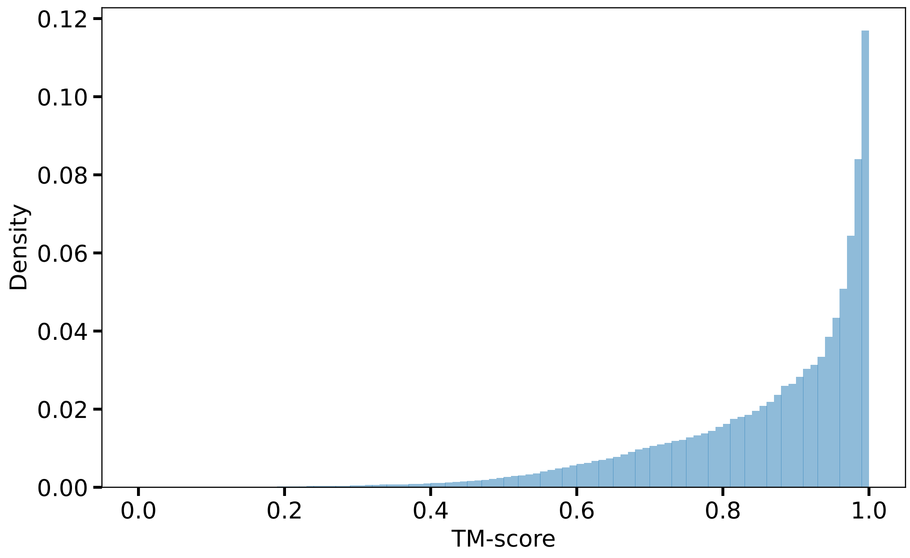


Figure S3: Distribution of pairwise structure comparison scores of proteins inside ECOD groups. Given an ECOD group with $N$ members, $N^2$ comparisons were attempted. This was repeated for all groups in ECOD. The figure shows distribution of structural similarity scores where comparisons were successful. The tail of the distribution extending towards 0 indicates low pairwise structural similarity between proteins in the same group.

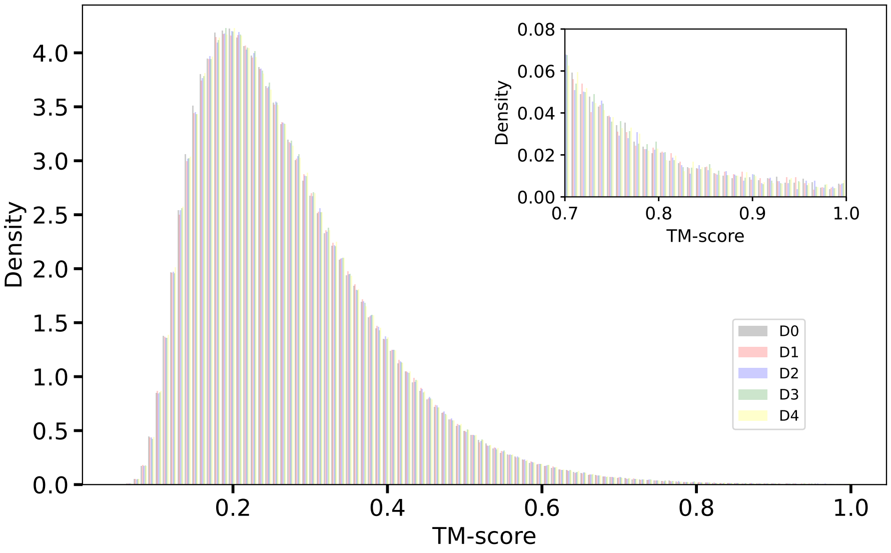


Figure S4: Distribution of pairwise structure comparison scores of proteins across SCOP groups. Given the computational complexity of comparing all protein structures in all groups to all other proteins in all other groups, five small datasets were created. Each dataset comprised one member randomly selected from a group as a representative of that group. The figure shows the distribution of pairwise comparison scores from the five datasets ($D0 - D4$). The figure shows distribution of structural similarity scores where comparisons were successful. The tail of the distribution extending towards 1 indicates high pairwise structural similarity between proteins across groups. Values of 1 arise from self comparison.

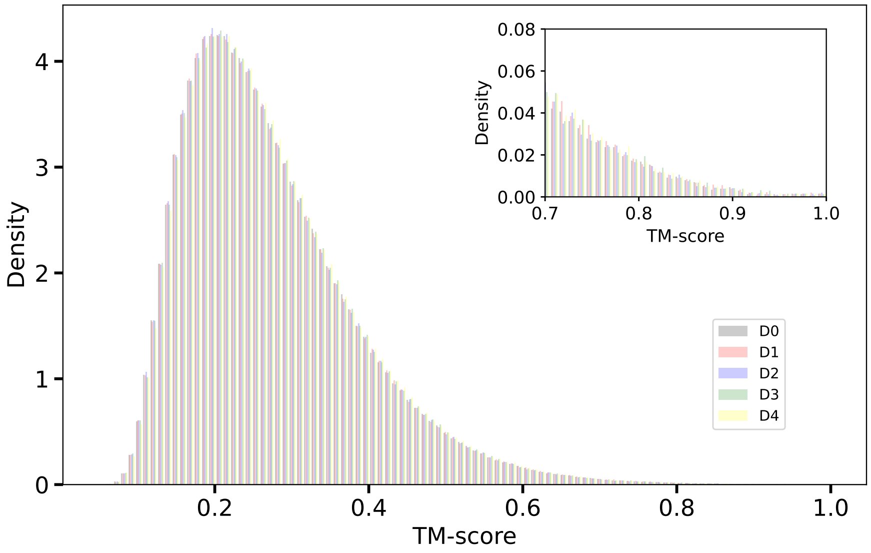


Figure S5: Distribution of pairwise structure comparison scores of proteins across CATH groups. Given the computational complexity of comparing all protein structures in all groups to all other proteins in all other groups, five small datasets were created. Each dataset comprised one member randomly selected from a group as a representative of that group. The figure shows the distribution of pairwise comparison scores from the five datasets (D0 − D4). The figure shows distribution of structural similarity scores where comparisons were successful. The tail of the distribution extending towards 1 indicates high pairwise structural similarity between proteins across groups. Values of 1 arise from self comparison.

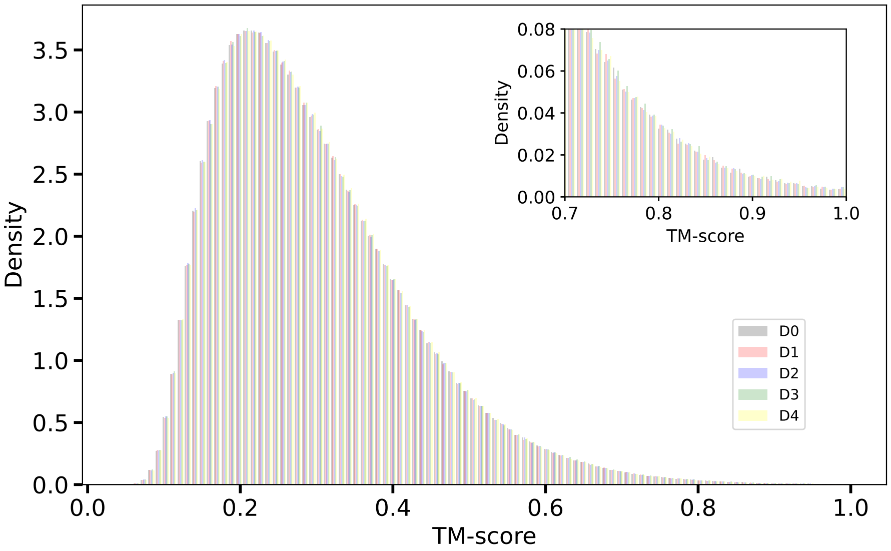


Figure S6: Distribution of pairwise structure comparison scores of proteins across ECOD groups. Given the computational complexity of comparison all protein structures in all groups to all other proteins in all other groups, five small datasets were created. Each dataset comprised one member randomly selected from a group as a representative of that group. The figure shows the distribution of pairwise comparison scores from the five datasets (D0 − D4). The figure shows distribution of structural similarity scores where comparisons were successful. The tail of the distribution extending towards 1 indicates high pairwise structural similarity between proteins across groups. Values of 1 arise from self comparison.

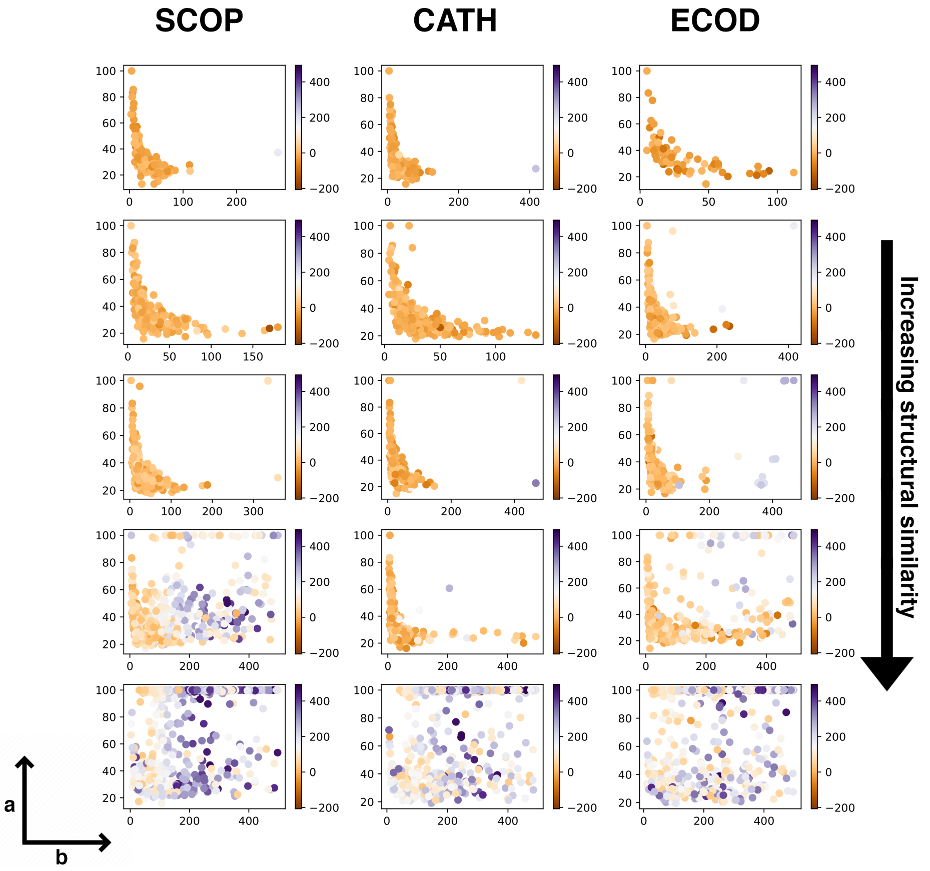


Figure S7: Trends in sequence-based metrics with increasing structural similarity. The figure grid shows comparison of protein pairs for SCOP, CATH and ECOD respectively with sequence similarity on (a) the vertical axis and (b) number of residues aligned in the sequence alignment on the horizontal axis for all plots. For a given comparison colour represents HMM scores. Compared to Figure 2, the signal obtained from structure comparison metrics is higher at the third and fourth levels of assignments (i.e., pairs with similar H1.H2.H3 [row 4] and H1.H2.H3.H4 [row 5], see methods for more details).

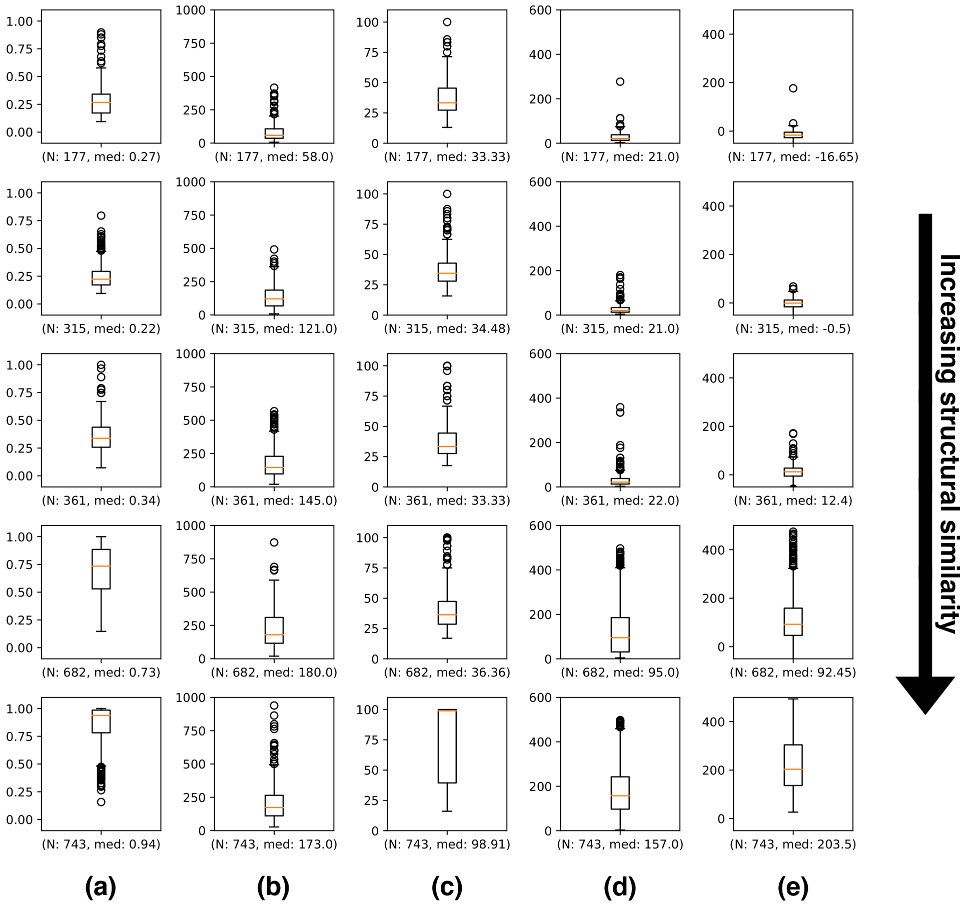


Figure S8: Distributions of comparative statistics for SCOP. The figure grid shows in five columns distributions of a) structural alignment score [0-1], b) residues aligned in structural alignment, c) sequence similarity score [0-100%], d) residues aligned in sequence alignment, e) average HMM score. The structural similarity increases along the rows from top to bottom. See methods for more details. Structure-bound metrics, (a,b,e) show progressive increments as degree of structural similarity increases. For each plot "$N$" is the number of data points constituting the distribution with "$med$" showing the median.

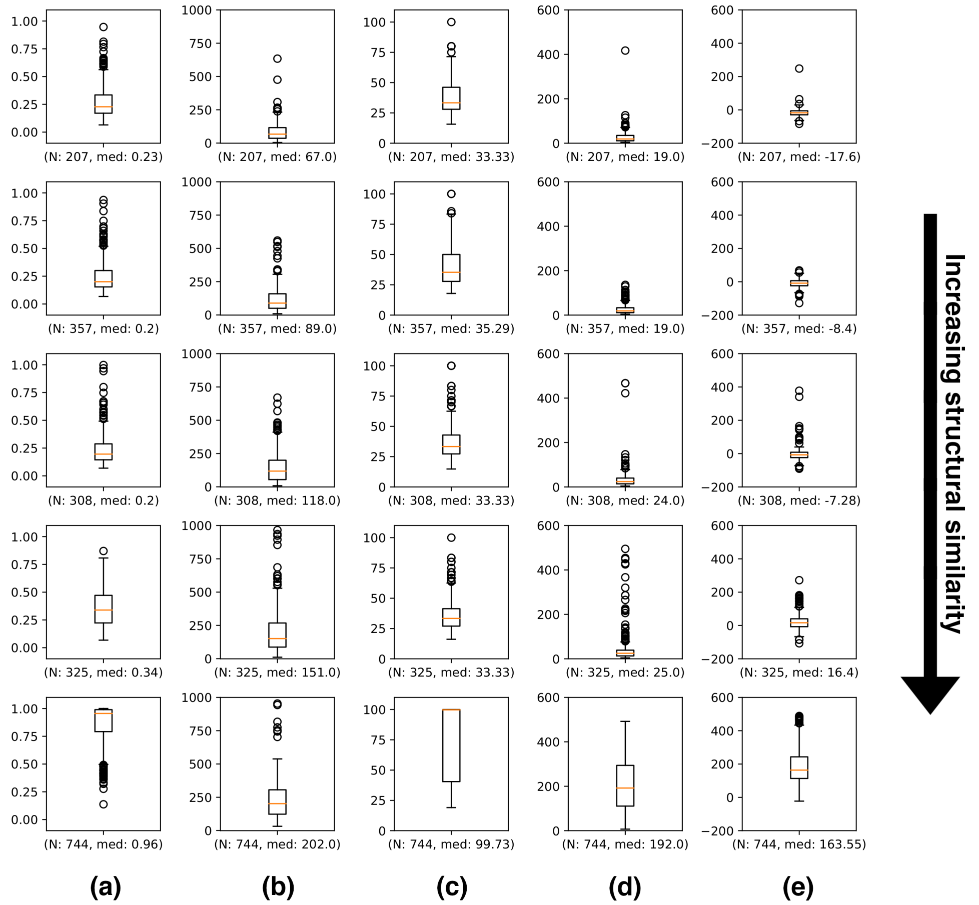


Figure S9: Distributions of comparative statistics for CATH. The figure grid shows in five columns distributions of a) structural alignment score [0-1], b) residues aligned in structural alignment, c) sequence similarity score [0-100%], d) residues aligned in sequence alignment, e) average HMM score. The structural similarity increases along the rows from top to bottom. See methods for more details. Structure-bound metrics, (a,b,e) show progressive increments as degree of structural similarity increases. For each plot "$N$" is the number of data points constituting the distribution with "$med$" showing the median.

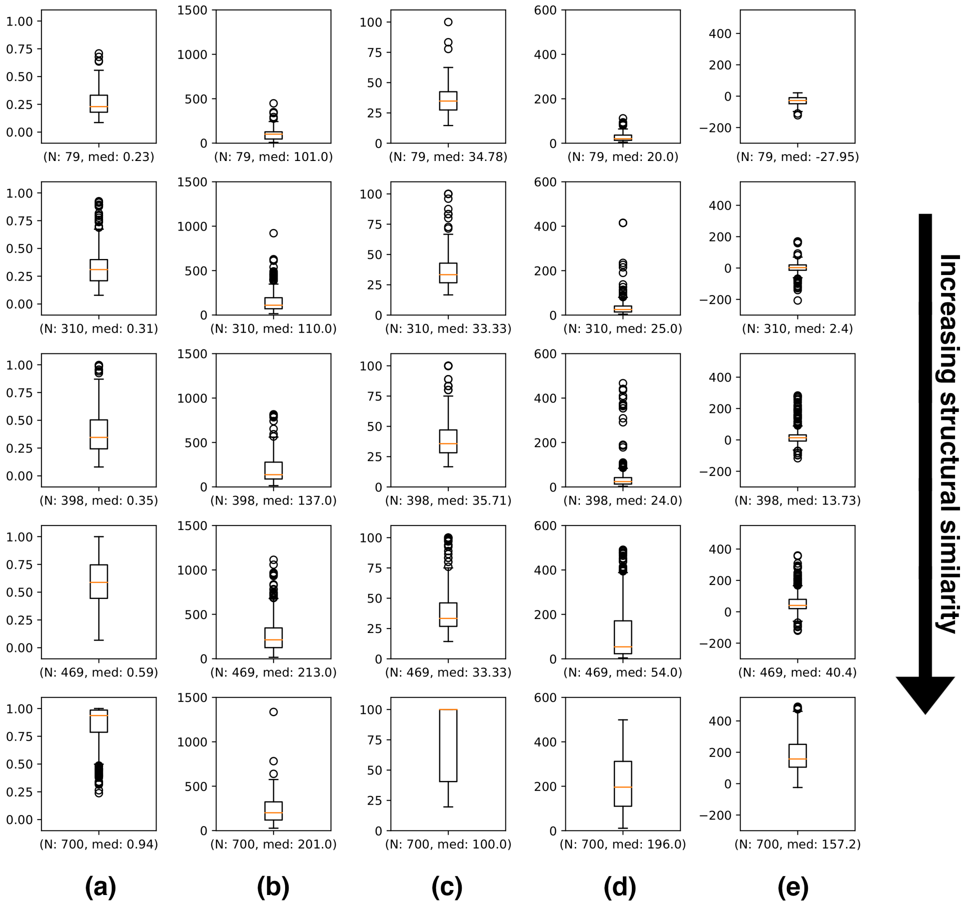


Figure S10: Distributions of comparative statistics for ECOD. The figure grid shows in five columns distributions of a) structural alignment score [0-1], b) residues aligned in structural alignment, c) sequence similarity score [0-100%], d) residues aligned in sequence alignment, e) average HMM score. The structural similarity increases along the rows from top to bottom. See methods for more details. Structure-bound metrics, (a,b,e) show progressive increments as degree of structural similarity increases. For each plot "$N$" is the number of data points constituting the distribution with "$med$" showing the median.

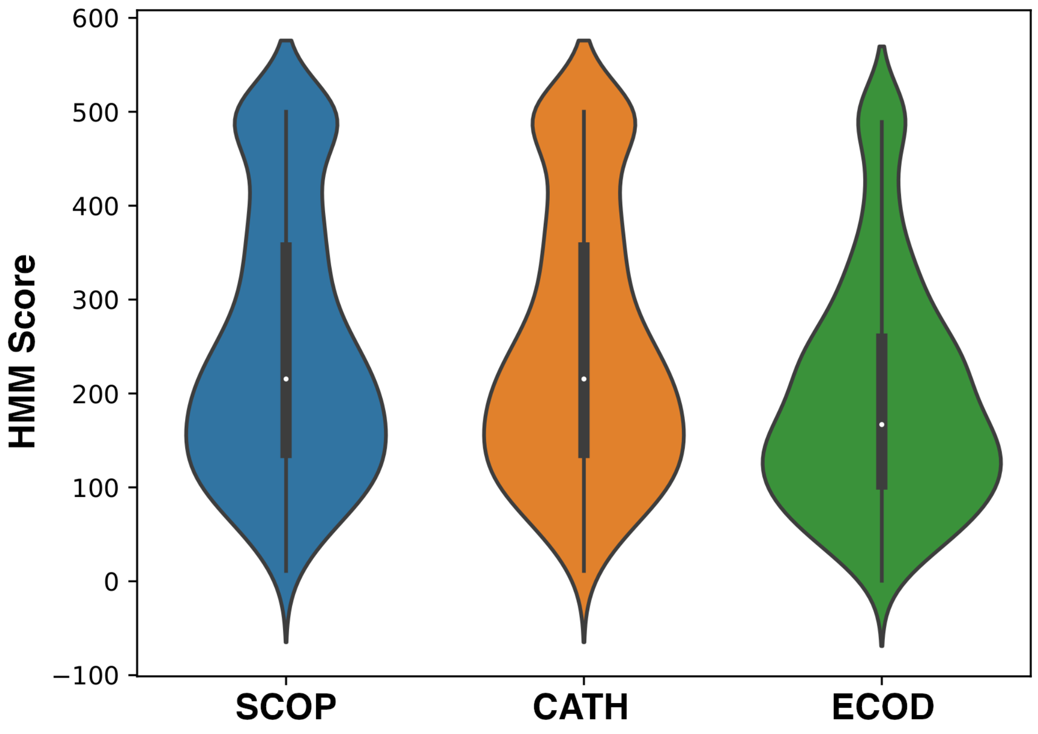


Figure S11: Comparison of proteins to their respective models. For a collection of $N$ proteins in a group $N-1$ proteins were used to develop the HMM model for that group. The excluded protein was then compared to its respective model. The distribution of HMM scores of >500 such comparisons for each SCOP, CATH and ECOD are shown, with higher HMM scores saturated at 600 for brevity. Plots show that proteins that belong to a group on average score highly against it.See supplementary table for data.

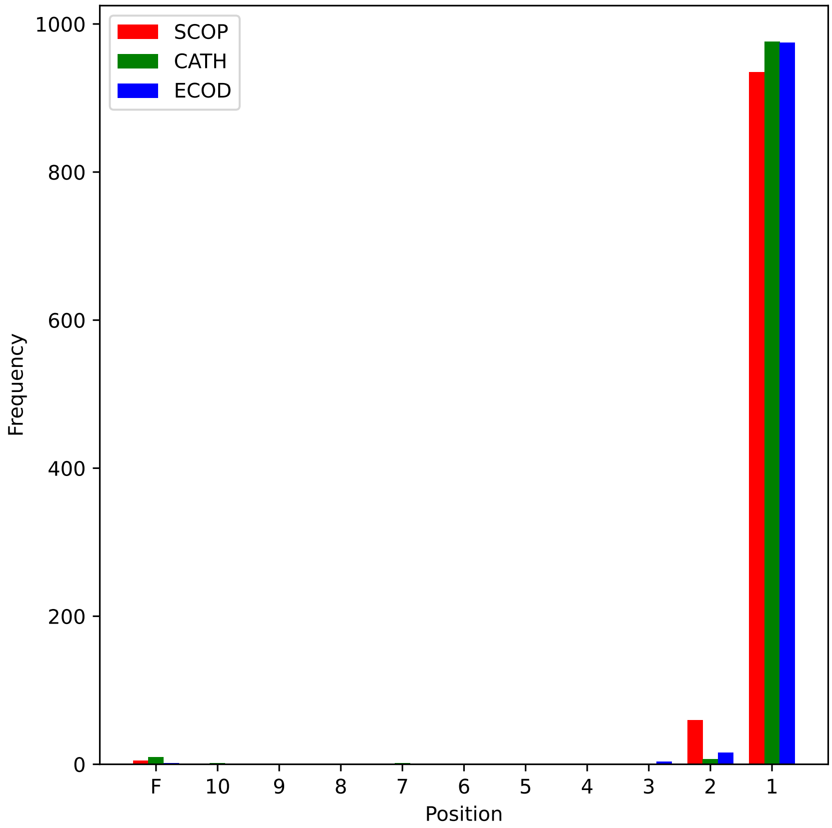


Figure S12: Position of true assignment in top-10 predicted results. A set of 1000 structures each for SCOP, CATH and ECOD, where the true grouping was known was used. The barplot presents the position of the true assignment in the top-10 predicted groupings. If the true grouping was not present in the top-10, a fail label "F" was assigned. The plot shows that for each SCOP, CATH and ECOD, the true results are mostly recovered in the top two hits.
